# Bayesian optimization for design of multiscale biological circuits

**DOI:** 10.1101/2023.02.02.526848

**Authors:** Charlotte Merzbacher, Oisin Mac Aodha, Diego A. Oyarzún

**Affiliations:** School of Informatics, University of Edinburgh, Edinburgh EH8 9AB, UK; The Alan Turing Institute, London, NW1 2DB, UK; School of Biological Sciences, University of Edinburgh, Edinburgh EH9 3JH, UK

**Keywords:** bayesian optimization, machine learning, dynamic pathway control, genetic circuit design, multiscale biological systems, metabolic engineering

## Abstract

Recent advances in synthetic biology have enabled the construction of molecular circuits that operate across multiple scales of cellular organization, such as gene regulation, signalling pathways and cellular metabolism. Computational optimization can effectively aid the design process, but current methods are generally unsuited for systems with multiple temporal or concentration scales, as these are slow to simulate due to their numerical stiffness. Here, we present a machine learning method for the efficient optimization of biological circuits across scales. The method relies on Bayesian Optimization, a technique commonly used to fine-tune deep neural networks, to learn the shape of a performance landscape and iteratively navigate the design space towards an optimal circuit. This strategy allows the joint optimization of both circuit architecture and parameters, and hence provides a feasible approach to solve a highly non-convex optimization problem in a mixed-integer input space. We illustrate the applicability of the method on several gene circuits for controlling biosynthetic pathways with strong nonlinearities, multiple interacting scales, and using various performance objectives. The method efficiently handles large multiscale problems and enables parametric sweeps to assess circuit robustness to perturbations, serving as an efficient *in silico* screening method prior to experimental implementation.

## 1 Introduction

The design of molecular circuits with prescribed functions is a core task in synthetic biology (*1*). These circuits can include components that operate across various scales of cellular organization, such as gene expression, signalling pathways (*2*) or metabolic processes (*3*). Computational methods are widely employed to discover circuits with specific dynamics (*4–6*) and, in particular, optimization-based strategies can be employed to search over design space and single out circuits predicted to fulfil a desired function (*7–10*). However, circuit design requires the specification of circuit architecture, i.e. the circuit “wiring diagram”, as well as the strength of interactions among molecular components. Since circuit architectures are discrete choices and molecular interactions depend on continuous parameters such as binding rate constants, circuit design leads to mixed-integer optimization problems that can be notoriously difficult to solve (*11*). Moreover, when circuits operate across multiple scales, their computational models become numerically stiff (*12*), resulting in extremely slow simulations that make their mixed-integer optimization challenging or even impossible to solve.

Previous work on computational circuit design has largely focused on genetic circuits that operate in isolation from other layers of the cellular machinery (Figure 1A). A range of techniques have been employed to identify functional circuits, including exhaustive search (*4–6, 13*), computational optimization (*7, 8*), systems theoretic approaches (*14–18*), Bayesian design (*19, 20*), and machine learning (*9, 21*). While these methods differ on their specific modelling strategies and assumptions, they all require computational simulations at many, typically thousands to millions, locations in the design space. But since multiscale systems often cannot be simulated at such scale, the computational costs limit the applicability of current optimization methods.

**Figure 1:**
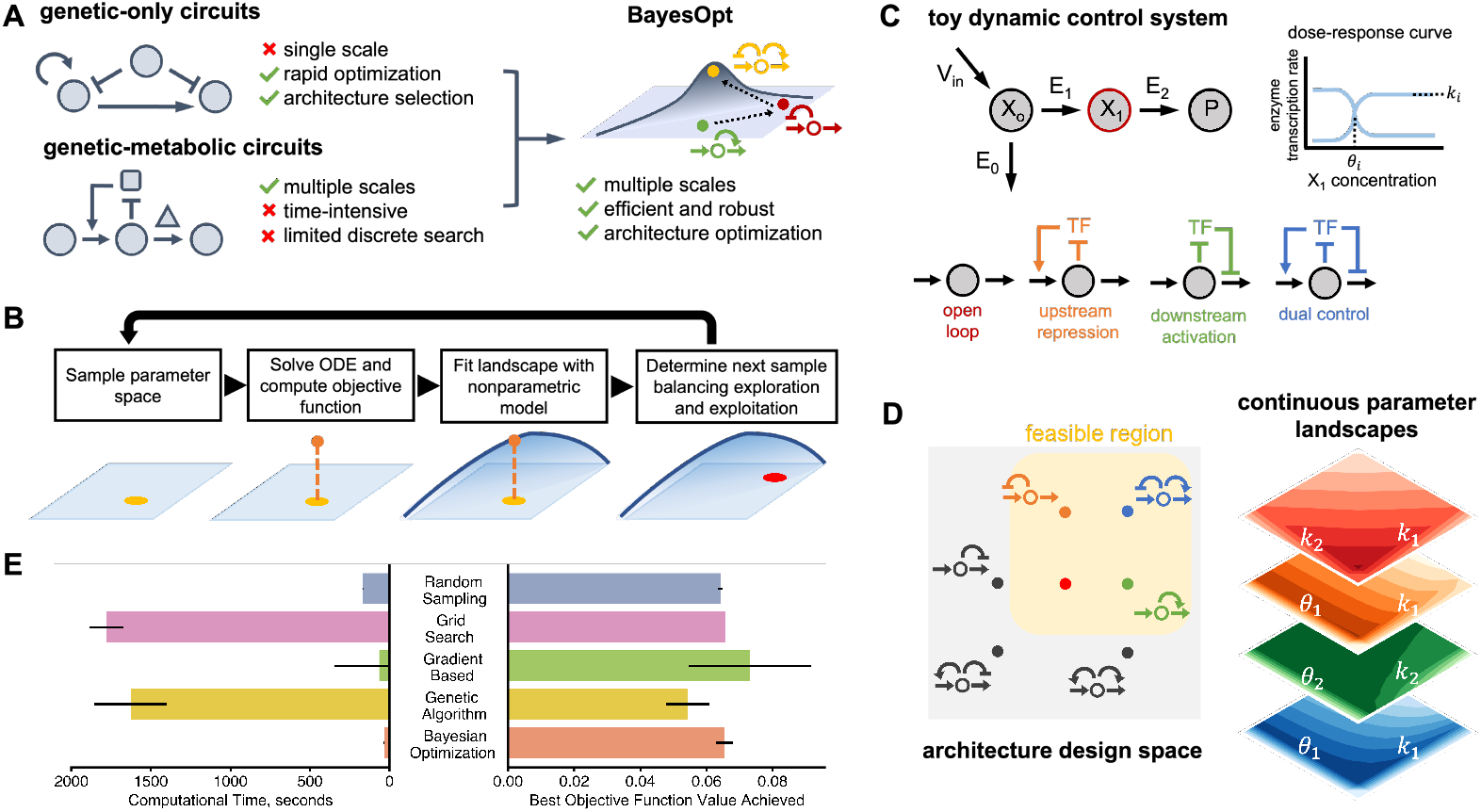
Bayesian optimization for the design of circuit architectures and parameters. **(A)** Previous optimization methods have focused on genetic circuits in isolation from other cellular processes. For multiscale circuits, optimization approaches become infeasible due to the difficulty of simulating stiff dynamical systems in many locations of the design space; a common example of such multiscale systems are gene circuits that control metabolic production (*3*). We propose the use of Bayesian optimization (BayesOpt) for efficient optimization of architectures and parameters in multiscale circuits. **(B)** Schematic of a mixed-integer Bayesian optimization loop; the objective function is regarded a random variable to be optimized over an input space comprised of continuous parameters and a set of discrete circuit architectures. At each iteration, the algorithm computes the value of the objective function from the solution of an ordinary differential equation (ODE) model at a single location in the input space. The algorithm learns the shape of the objective landscape using a nonparametric statistical model (*37*), which is employed to propose a new location in the input space through an “acquisition function” designed to balance exploration and exploitation of the input space; more details in Methods. The algorithm iteratively learns the shape of the performance landscape until convergence to a global optimum. **(C)** Example metabolic pathway under gene regulation. We consider three negative feedback architectures plus open loop control; the architectures are named based on the net effect of the metabolite on gene expression. The intermediate *X*_1_ binds a transcription factor (TF) that controls the expression of pathway enzymes, either as an activator or repressor. *V*_in_ is the constant influx to the engineered pathway from native metabolism. The TF dose-response curve (at right) is described by three parameters, *k*_*i*_, *θ*_*i*_, and *n*, where *i* = 1, 2. The aim is to find designs with optimal architecture and dose-response parameters (*k*_*i*_, *θ*_*i*_); for simplicity the Hill coefficient was fixed to *n*_*i*_ = 2. **(D)** Performance landscapes of the four feasible circuit architectures. We exclude architectures with positive feedback loops as these are prone to multistability (*47*). The shape of the performance landscape defined in Eq. (3) shows substantial variation across the four architectures. This leads to a highly non-convex mixed-integer optimization problem. Heatmaps show the value of the objective *J* computed on a regular grid of the indicated parameters. **(E)** Comparison of BayesOpt against other strategies using the toy model as a benchmark; lower objective function values are better. Shown are the results for random sampling (*N* = 1, 000 samples), grid search (*N* = 40, 000), a genetic algorithm (*55*) (*N* = 100 individuals, *N* = 1000 generations), and a gradient-based optimizer to find optimal continuous parameter values for each architecture (*48*).

A notable example of this challenge appears in genetic circuits for dynamic control of metabolic pathways (*22–26*). These systems are receiving substantial attention thanks to several successful implementations that improved yields as compared to classic techniques in metabolic engineering (*27, 28*). The key principle is to put enzymatic genes under the control of metabolite-responsive mechanisms that couple heterologous expression to the concentration of a pathway intermediate (*3*). This creates feedback loops between enzyme expression and pathway intermediates that allow controlling pathway activity in response to upstream changes in growth conditions or precursor availability. Such dual genetic-metabolic systems are particularly challenging to simulate efficiently because metabolites and enzymes vary in different timescales, from milliseconds (enzyme kinetics) to minutes (enzyme expression), and they also appear in vastly different concentrations; in bacteria enzymes are typically expressed in nanomolar concentrations, whilst metabolites are found typically above the millimolar range (*29*). Moreover, the implementation of these systems is costly and requires substantial experimental fine-tuning. As a result, a central task prior to implementation is the choice of a suitable feedback control loops between metabolites and enzymatic genes, and the strength of interactions between metabolites and actuators of gene expression such as transcription factors (*30*) or riboregulators (*31*). The design of control architectures is particularly important, because there are many ways of building similar control loops (*32*), for example by employing combinations of transcriptional activators and repressors (*33, 34*), that may differ in their performance and cost of implementation.

Here, we present a fast and scalable machine learning approach for optimization of multiscale circuit architectures and parameters (Figure 1A). The method is based on Bayesian optimization coupled with differential equation models, and we highlight its utility in various models of metabolic pathways under genetic feedback control (*35*). Using a toy example for a simple pathway, we first show that the method converges rapidly and outperforms other optimizers by a substantial margin. We then consider real-world models of metabolic pathways in *Escherichia coli* for the production of several relevant precursors: glucaric acid (*36*), fatty acids (*33*), and p-aminostyrene (*34*). We use these pathways to illustrate how the speed of our method enables screening optimal designs in realistic design tasks that would otherwise be infeasible to compute, including the impact of uncertain enzyme kinetic parameters, the use of layered architectures that combine metabolic and genetic control, and the optimization of a complex model with 23 differential equations, 27 candidate control architectures, and 19 parameters to be optimized. The method can help speed up the design of synthetic biological circuits and presents a novel approach to explore the design space ahead of implementation.

## 2 Results

### 2.1 Bayesian optimization for joint optimization of circuit architecture and parameters

In general, a circuit design task can be stated as the following mixed-integer optimization problem:

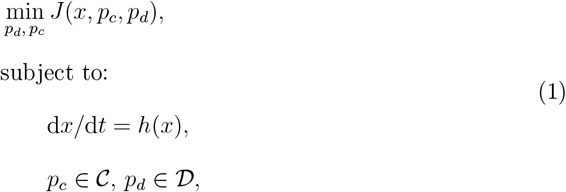

where *J*(*x, p*_*c*_, *p*_*d*_) is a performance objective to be optimized over a space of continuous parameters *p*_*c*_ and a discrete set of circuit architectures *p*_*d*_. The ODE in Eq. (1) describes the temporal dynamics of circuit components and are typically built from mass balance relations comprised in the nonlinear function *h*(*x*). Common examples of continuous parameters in applications are binding affinities between DNA and regulatory proteins, or the strength of protein-protein interactions. Conversely, circuit architecture would typically involve various combinations of positive and negative feedback loops among molecular species. We have stated the problem as minimization of *J*, but similar formulations can be posed as a maximization problem.

In this paper, we propose to solve the design problem in Eq. (1) with Bayesian optimization (BayesOpt), a class of algorithms designed for problems with objective functions that are expensive to compute. BayesOpt is a global optimization technique that treats the objective function as a random variable with a prior distribution on it. The algorithm creates a statistical model of the objective through subsequent evaluations, which are employed to build a posterior distribution and determine the next set of inputs to evaluate (Figure 1B). A typical application of BayesOpt is in design of experiments (*35*) where the objective function requires measuring data with costly and/or slow experimental work. In deep learning, BayesOpt is widely employed for model selection, as traditional grid search approaches require large compute resources to train many architectures with combinations of various layer sizes and other hyperparameters (*37, 38*).

For the circuit design task in Eq. (1), if the biological system has multiple scales the computation of objective *J* requires solving a stiff ODE in many locations of the mixed-integer search space, which can rapidly become infeasible. To illustrate the utility of BayesOpt in a range of design problems, we focus on genetic control circuits for metabolic pathways that synthesize high-value products. In this case, the ODE in Eq. (1) contains two sets of equations:

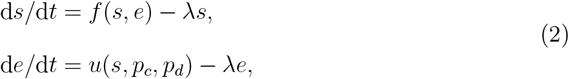

where *s* and *e* are vectors of metabolite and enzyme concentrations, respectively. The term *f*(*s, e*) describes the biochemical reactions between pathway intermediates, while the parameter *λ* models the dilution effect by cell growth. The vector *u*(*s, p*_*c*_, *p*_*d*_) describes the enzyme expression rates controlled by some pathway intermediates, and typically take the form of sigmoidal dose-response curves that lump together processes such as metabolite-TF or metabolite-riboregulator interactions (*30*). The continuous parameters *p*_*c*_ model the dose-response curves of the feedback mechanisms, whereas the discrete parameters *p*_*d*_ specify the gene control architecture. The number of heterologous enzymes determines the number of genes in the control circuit. In pathways under dynamic control as in Eq. (2), both sets of species change in vastly different timescales; metabolic reactions operate in the millisecond range or faster (*39*), whilst enzyme expression changes in the scale of minutes or longer. Moreover, metabolites and enzymes are also present in different ranges of concentrations, from nM for enzymes to mM and higher for metabolites (*29*). As a result, simulation of the ODE in Eq. (2) is computationally expensive, particularly when this has to be done many times as part of an optimization-based search.

The performance objective *J* can be flexibly used to model common design goals such as production flux, yield or titer, as well as cost-benefit tasks that balance production with the deleterious impact of the pathway on the physiology of the host. To first establish a baseline for the performance of our method, we employed a simple toy pathway model that displays common features found in real metabolic pathway (Figure 1C). The model includes a metabolic branch point through a heterologous pathway with two enzymatic steps. As a performance objective we considered the minimization of

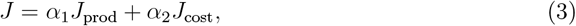

where *J*_prod_ was designed so that its minimization is equivalent to maximization of the production flux, and *J*_cost_ penalizes total amount of enzyme expressed during the culture. The parameters *α*_1_ and *α*_2_ are positive weights used to control the balance between the costs and benefits of expressing the heterologous pathway. Details on the objective function can be found in the Supplementary Information.

We considered the four control architectures shown in Figure 1C, which include open loop control as well as three different implementations of negative feedback control using a metabolite-responsive transcription factor. Negative feedback is widely employed in gene circuits as it has substantial benefits in terms of robustness and performance, and their properties have been extensively studied in the literature (*40–42*). To illustrate the challenge of jointly optimizing circuit architecture and parameters, in Figure 1D we show a schematic of the design space. The four control architectures under consideration reside at different discrete points in the architecture space. Within each architecture, we observe substantial variations in the shape of the performance landscape *J* as a function of the dose-response parameters *p*_*c*_. There are cases with convex landscapes with a clear optimum (e.g. dual control) and landscapes with flat basins where most optimization algorithms would struggle to find the optimum (e.g. downstream activation). When searching over the space of architectures and parameters simultaneously, the problem becomes a mixed-integer, non-convex optimization that is extremely challenging to solve with traditional approaches.

We implemented a BayesOpt routine to jointly compute the architecture (*p*_*d*_) and dose-response parameters (*p*_*c*_) that minimize the performance objective in Eq. (3). We bench-marked its performance against several other methods, including a random search, an exhaustive grid search, a gradient based method, and a genetic algorithm (Figure 1E). The algorithm was able to compute optimal solutions rapidly (average 27 seconds per run across 100 runs) and robustly (standard deviation less than 2.5% of the mean optimal objective function value). BayesOpt runs significantly faster than the other methods, and provides a 30-fold improvement over a genetic algorithm. The accuracy of the optimum, quantified by the minimal value of the objective function, is on average 11.4% worse than the genetic algorithm, but this falls within the variation of the latter across several runs. We also note that the traditional gradient-based optimizer proved unreliable and failed to converge on 14.5% of runs.

One key advantage of Bayesian methods is that they are not gradient-based, and therefore are not constrained to navigate the space smoothly in the direction of steepest descent. Gradient-based methods can get trapped in local minima and struggle to find the global optimum, especially in highly nonconvex landscapes like the ones presented here. In contrast, BayesOpt does not converge by chasing minima directly but rather by modelling the entire objective function landscape, which results in rapid and reliable results. The method can perform multiple “jumps” between distant locations in the discrete-continuous search space, where each subsequent sample is selected to maximize the expected improvement on the best sample found so far.

The speed of our approach enables the computation of large solution ensembles under model perturbations such as sweeps of key model parameters. In addition, our method can search high-dimensional mixed-integer design spaces. We next illustrate the versatility of the approach in a range of relevant real-world pathways that require solving the optimization problem for large samples of parameter values.

### 2.2 Robustness of control circuits to uncertainty in enzyme kinetic parameters

A challenge in building pathway models is the substantial uncertainty on the enzyme kinetic parameters; this is particularly critical for pathways that include regulatory mechanisms such allostery or product inhibition, which are often poorly characterized. Databases such as BRENDA (*43*) often have insufficient data on enzyme kinetics for a particular host strain or substrate of interest. Since pathway dynamics can strongly depend on enzyme kinetics, the parametric uncertainty requires extensive sweeps of kinetic parameters to determine the robustness of a specific control architecture deemed to be optimal.

We focused on a pathway for synthesis of glucaric acid in *E. coli* (Figure 2A), a key precursor for many downstream products (*36*). The pathway branches from glucose-6-phosphate (g6p) in upper glycolysis and contains three enzymatic steps (Ino1, SuhB, and MIOX). Doong and colleagues implemented a dynamic control circuit using the dual transcriptional regulator IpsA which responds to the intermediate myo-inositol (MI) (*26*). The pathway enzyme MIOX is allosterically activated by its own precursor, and one intermediate MI, can be exported to the extracellular space. We employed a previously developed ODE model (*10*) that was parameterized using a combination of enzyme kinetic data and omics measurements, and considered the same four control architectures as in the previous example, including various alternative implementations of negative feedback control.

**Figure 2:**
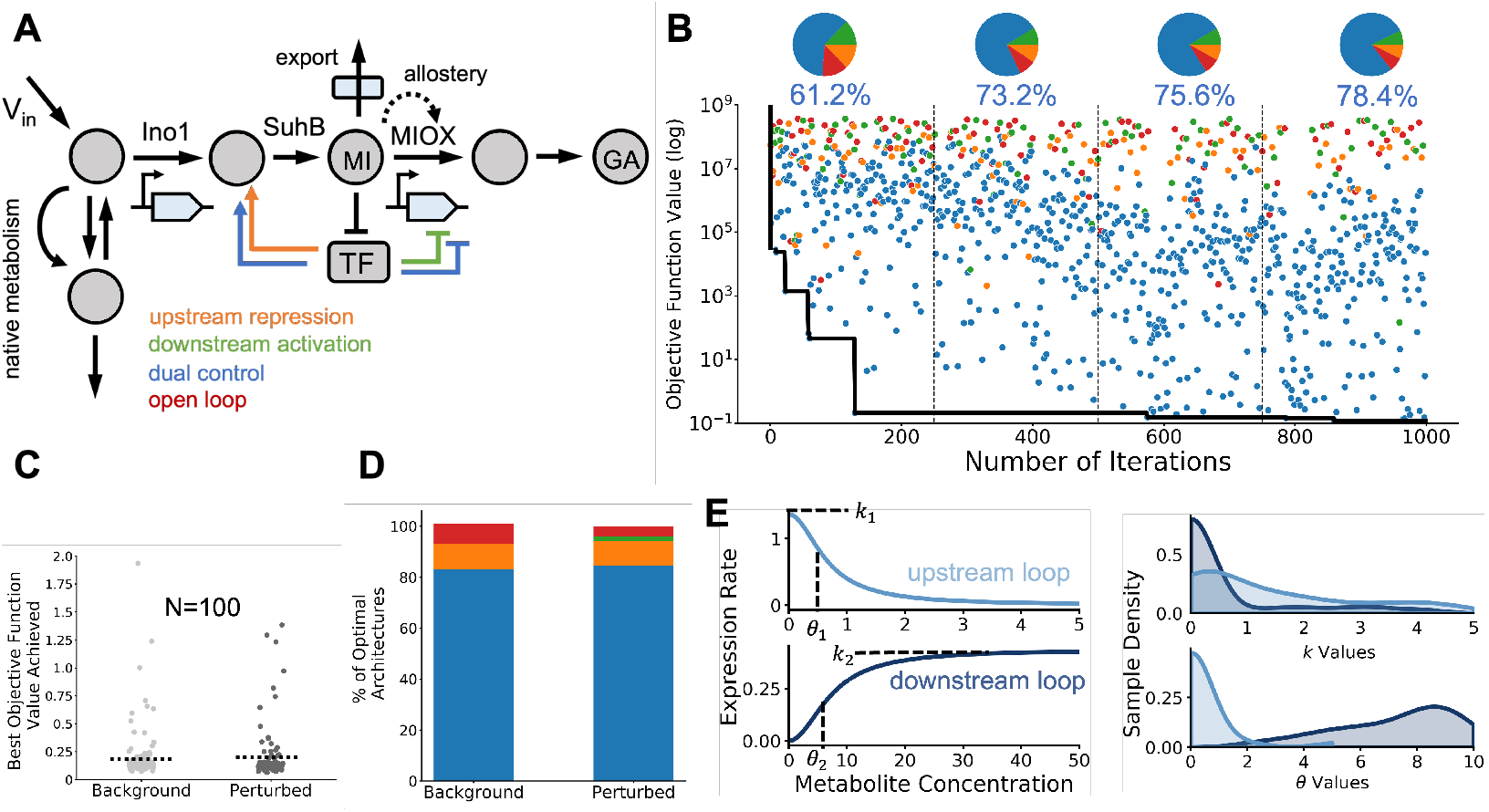
Robustness optimal circuits to parameter uncertainty. **(A)** Schematic of a dynamic pathway for production of glucaric acid in *Escherichia coli* (*26*). The pathway includes allosteric inhibition and export of an intermediate to the extracellular space. The core pathway components myoinositol (MI) and glucaric acid (GA) are modelled explicitly, as are the enzymes Ino1 and MIOX. The enzyme SuhB is not rate-limiting and is not modeled explicitly. *V*_in_ is the constant influx to the engineered pathway from native metabolism. As in Figure 1C, the architectures are named based on the net effect of the metabolite on gene expression. **(B)** Sample run of the BayesOpt algorithm for 1,000 iterations of the loop in Figure 1B. Black line shows the descent on the value of the objective function. Dots show all samples colored by architecture; pie charts show the fraction of architectures explored by the algorithm, and the fraction of samples taken from the majority architecture (dual control). The first quarter of the run had the most exploration of architectures other than dual control, with 38.6% of samples coming from non-majority architectures. This percentage steadily decreased over the iterations but did not drop below 20%, illustrating the global nature of the optimization routine. **(C)** To examine the robustness of the optimal solutions to parameter uncertainty, we computed optimal solutions for many perturbed parameters of the allosteric activation of MIOX by its substrate myo-inositol (MI). Strip plot shows the best objective function values achieved for background and perturbed kinetic parameters (*V*_m,MIOX_, *a*_MIOX_, *k*_a,MIOX_) in Eq. (4). Kinetic parameters were perturbed using Latin Hypercube sampling (*56*) on the range (-100%, +100%) of the nominal values (Supplementary Information). We observed little difference between between background and perturbed values; dashed line denotes the mean value of the objective function. Only one of the *N* = 100 runs for perturbed parameters failed to converge the optimum. **(D)** Optimal architectures across runs with background and perturbed parameter values. Both background and perturbed systems resulted in over 80% of runs selecting dual control as the optimal architecture. **(E)** Average dose-response curves and distribution of optimal parameters for the dual control architecture with perturbed allosteric parameters. The repressive and activatory loops have substantially different dose-response curves on average. The distributions of the dose-response parameters (right) show important variations in their mean and dispersion. The parameter *k*_*i*_ and *θ*_*i*_ determine the maximal enzyme expression rate and regulatory threshold, respectively.

The results in Figure 2B show a typical run of the optimizer when using the cost-benefit objective in Eq. (3) (details in Supplementary Information), together with the fraction of samples in which the algorithm explored each control architecture across the successive iterations. The optimal architecture (dual control in this case) was found quickly and the algorithm was able to further decrease the value of the objective function by exploring the space of dose-response parameters of IpsA. We observe that as the iterations progress, the algorithm shows a remarkable ability to explore other architectures despite their larger objective function values, thus highlighting the global nature of the algorithm.

To explore the impact of uncertain enzyme kinetics, we perturbed the parameters of the rate-limiting MIOX allosteric reaction:

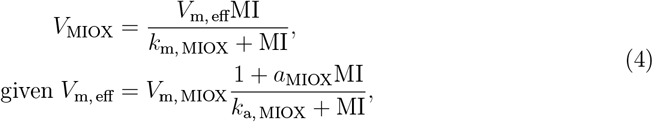

where *V*_m, MIOX_ is the maximum rate of reaction, *k*_m, MIOX_ is the Michaelis-Menten constant, and *k*_a, MIOX_ and *a*_MIOX_ are allosteric activation constants. We solved the optimization problem for 1000 combinations of these three parameters, which took under 16 hours on a Mac-book Air with Apple M1 processor and 8GB of RAM running MacOS Monterey. Perturbing the kinetic parameters of the glucaric acid pathway did not significantly affect the minimum objective function value achieved, indicating that the optimum is robust to uncertainty in the kinetic parameters (Figure 2C). However, the mean optimal objective function value was not significantly higher among the perturbed samples. We found that the dual control architecture was chosen as optimal in more than 85% of samples (Figure 2D). We thus sought to examine the optimal dose-response parameters of this architecture in more detail.

The maximal enzyme expression rates (*k*) and regulatory thresholds (*θ*) control the shape of the dose-response curves. As shown in Figure 2E, we found that the upstream repressive loop and downstream activatory loop had different optimal dose-response curves, corresponding to different optimal values of the continuous parameters. Optimal values of the upstream repression threshold *θ*_1_ are low (mean value 0.64) and compressed into a narrow range as compared to the larger standard deviation of the downstream repression threshold *θ*_2_ (mean value 7.24). This is reflected on a larger variation in the shape of the dose response curve for the downstream loop. Experimental fine-tuning of a dual control circuit might target parameters with optimal values with a wide range, such as *k*_1_, as varying these parameters is less likely to impair circuit function. Overall, these results show the robustness of the glucaric acid dual control system to kinetic parameter uncertainty and demonstrate the possibilities enabled by the speed of BayesOpt.

### 2.3 Exploration of alternative objective functions

In the previous case studies we employed a cost-benefit objective designed to account for the tradeoff between heterologous production and the cost of expressing pathway enzymes, as in Eq. 3. To demonstrate the flexibility of the method with other objective functions, here we consider the optimization of the temporal trajectories of pathway metabolites.

We focussed on the joint optimization of the rise time and overshoot in a model of a fatty acid production pathway considered previously in the literature (*33*). Fatty acids are an essential energy source and cellular membrane component. In addition, hydrocarbons derived from fatty acids have attracted attention as a potential biofuel source (*24, 44*). Recent work engineering metabolic and genetic control loops showed that negative feedback control could speed up the rise to steady-state conditions (*33*). The pathway built in literature expressed a thioesterase under transcriptional control, shown as the negative metabolic loop (NML) architecture in Figure 3A. In addition to transcription-factor mediated negative feedback loops, this model also includes individually implemented direct genetic loops where a repressor is expressed on the same promoter as the enzyme. These two different scales of loops interface with different levels of cellular organization. We explore several control architectures previously proposed in the literature (*33*) (Figure 3A).

**Figure 3:**
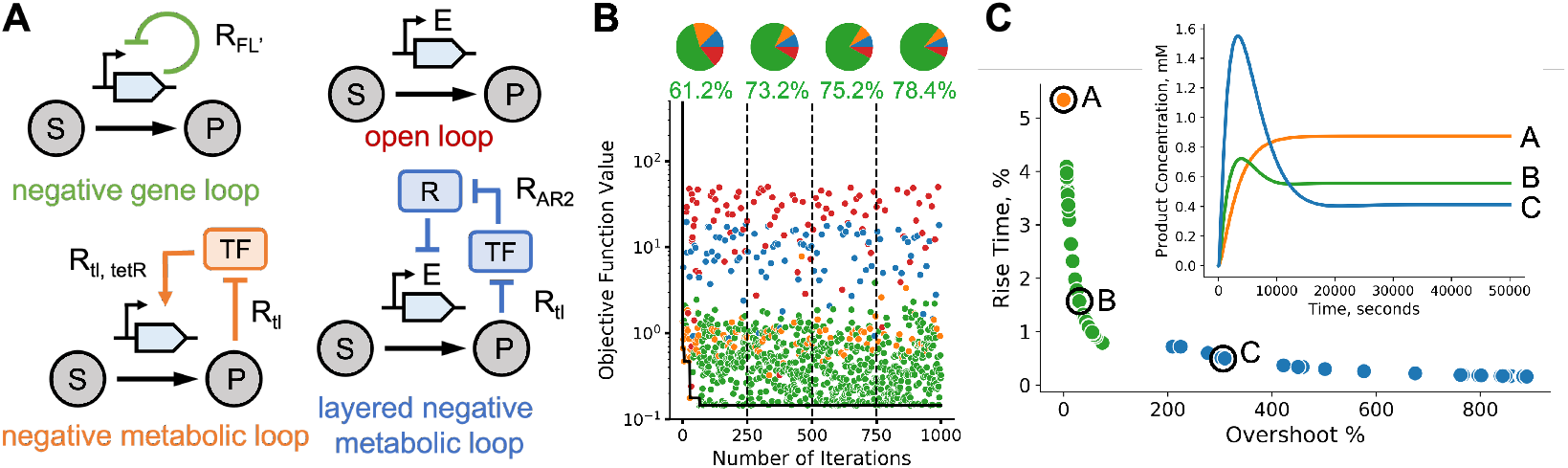
Performance landscapes of fatty acid synthesis pathway. **(A)** Pathway diagrams with various control architectures implemented in *Escherichia coli* (*33*). The metabolic loop employs a metabolite-responsive transcription factor, whereas the gene loop includes only a repressor expressed on the same promoter as the enzyme. **(B)** Representative run of BayesOpt with cost-benefit objective showing the best objective function value (black line). All samples are colored by their architecture. Pie charts of each quarter of the run show continued exploration of all architectures despite clear stratification in losses. **(C)** Optimal tradeoff curve between overshoot and rise time. The objective weight *α* was swept from *α* = 0.01 to *α* = 10, 000 and BayesOpt was run for 100 iterations at each *α* value. The optimal parameter values were used to compute the rise time and overshoot for visualization. The inset shows three sample trajectories illustrating how different optimal architectures navigate the tradeoff between overshoot and rise time.

We first considered a similar objective function as in Eq. (3) so as to compare convergence against the previous case studies. We implemented a modified production flux objective which takes the reciprocal of the product flux to convert the optimization to a minimization problem. The pathway cost *J*_cost_ is measured by summing the expression of all heterologous enzymes and varies across the different architectures. Details on the objective function can be found in the Supplementary Information. A representative optimization run for this objective (Figure 3B) shows that the negative gene loop (NGL, green) and negative metabolic loop (NML, orange) architectures perform, on average, better than the other two architectures. BayesOpt samples taken from the open-loop architecture were, on average, two orders of magnitude worse than samples taken from NML and NGL architectures. Despite such hierarchy of loss values across the four architectures, the method effectively explores all architectures throughout the optimization run.

We next considered the optimization of percent overshoot and rise-time presented in the literature (*33*). The percent overshoot objective, *J*_os_, measures the maximal deviation of product from its steady state concentration and is defined as the percent difference between the maximum fatty acid concentration and the steady-state fatty acid concentration. The rise-time, *J*_rt_, is a measure of how fast fatty acid production rises to steady-state and is defined as the first time point where the fatty acid concentration reaches 50% of the steadystate value, normalized by the total integration time. We minimized the sum of the overshoot and rise time with a scaling weight *α*:

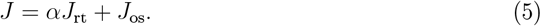

Adjusting *α* balances the relative importance of the two optimization criteria; details on the calculation of rise time and overshoot are in the Supplementary Information. Higher values of *α* correspond to optimal circuits with low rise-times, while lower values of *α* prioritize circuits with low overshoot. Rise-time is a measure of circuit speed, while overshoot is a measure of circuit accuracy. We found that when *α* is varied across several orders of magnitude, the optimal circuits form a optimal tradeoff curve (Figure 3C). Different architectures occupy different parts of the optimal tradeoff curve and display markedly different dynamics. The NML optima occupies a single point in the loss space, indicating that multiple continuous parameter values give the same loss function value for multiple values of *α*. The NML also has the lowest absolute loss function value of all the architectures considered. The NGL and layered negative metabolic loop (LNML) architectures occupy larger ranges on the curve, with NGL giving a low overshoot and LNML a low rise time. The optimal NML circuit has no overshoot but the slowest rise-time, while the LNML has a very rapid rise-time but overshoots the steady-state value by more than double. These opposing tradeoffs demonstrate the importance of balancing multiple circuit design objectives.

### 2.4 Scalability to large pathway models

Our previous case studies have been limited to circuits with a single metabolite controlling gene expression and a relatively small number of control architectures. We now study a large model for the synthesis of p-aminostyrene (p-AS), an industrially relevant vinyl aromatic monomer, in *E. coli* (Figure 4A) (*45*) using a cost-benefit objective similar to Eq. (3) tailored to the specific pathway (details in Supplementary Information). This model has two possible metabolites that can regulate gene expression, namely p-amminocinnamic acid (p-ACA) and p-aminophenylalanine (p-AF), both of which can act as ligands for aptazyme-regulated expression device (aRED) transcription factors (*46*), and three genes to be controlled. The aRED transcription factors can also act as dual regulators (activators or repressors) on any of the three promoters involved in the pathway. For simplicity, we limit the design space to control architectures without positive feedback loops, as these are prone to bistability (*47*). This results in 27 possible control architectures and 19 continuous parameters to be optimized. The model also has a number of additional complexities. It contains operon-based gene expression commonly found in bacterial systems (genes papA, papB, and papC are expressed on the papABC operon), it includes a detailed description of mRNA dynamics and protein folding, which results in a large model with 23 differential equations, and it can also display oscillatory dynamics.

**Figure 4:**
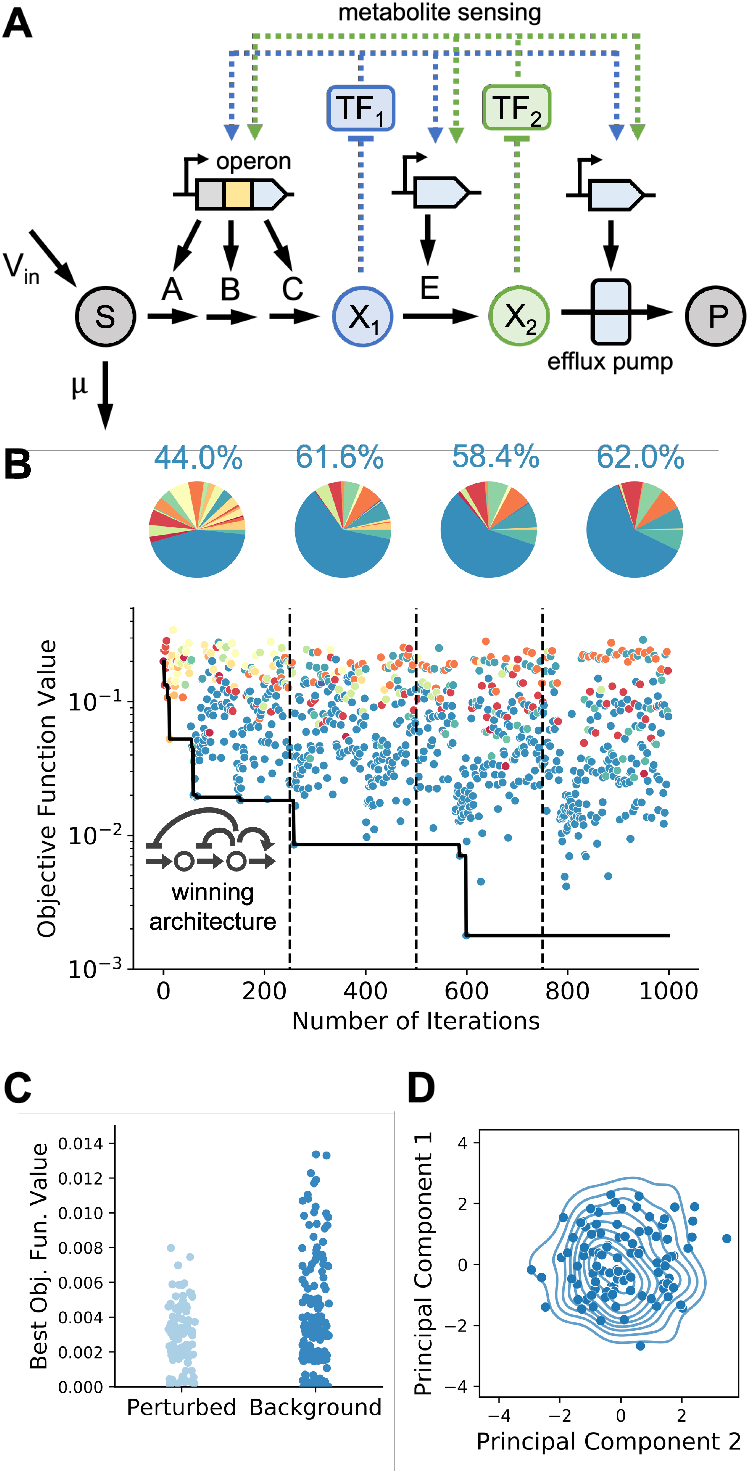
Bayesian optimization in a complex pathway. **(A)** Schematic of pathway for production of p-aminostyrene (*34*). Two intermediates can act as ligands for metabolite-dependent riboregulators, and three promoter sites of control. The optimization problem has 16 continuous decision variables and 27 circuit architectures. The substrate *S* is converted by enzymes *A, B*, and *C* to *X*_1_, which is then converted by *E* to *X*_2_. The toxic substrate *X*_2_ is then pumped out of the cell via an efflux pump to form the product *P*. Both *X*_1_ and *X*_2_ can act on the transcription factors TF_1_ and TF_2_. *V*_in_ is the constant influx to the engineered pathway from native metabolism. **(B)** Representative run of the BayesOpt algorithm; the method samples many architectures before settling on the optimal one. Pie charts show continued exploration of a large number of architectures. The winning architecture is shown in the inset. **(C)** The p-aminostyrene pathway has several forms of substrate, protein, and enzyme toxicity expressed via a toxicity factor *τ* (see Eq. (6)). To explore the effects of protein and metabolite toxicity, we perturbed the toxicity factor. Metabolite-induced toxicity was perturbed on the nominal range (1 ·10^−3^, 1 · 10^−4^) and protein-induced toxicity on the range (1 · 10^−4^, 1 · 10^1^) respectively. Both ranges were selected to match the ranges provided in the literature (*34*). Latin Hypercube sampling was used to generate *N* = 100 perturbed parameter values, and the optimal solutions were compared to an equal number of background solutions using the nominal parameter values. **(D)** Visualization of the optimal solutions; scatter plot of principal components of the optimal parameter values for the model with perturbed toxicity parameters (*N*=100). Contour plots show the background distribution of parameter values.

In addition to expression of heterologous enzymes, the accumulation of toxic intermediates is another major source of genetic burden to host organisms. The p-AS model has several sources of toxicity present in the pathway (*34, 45*). The intermediate p-ACA and the efflux pump used to remove p-ACA from cells are both cytotoxic, while another intermediate, p-AF, leaks from cells. The pathway enzyme L-Amino Acid Oxidase (LAAO) depletes key aromatic amino acid metabolites and creates toxic hydrogen peroxide as a byproduct. The model incorporates these various types of toxicity in the form of a toxicity factor tau. This toxicity factor is of the form

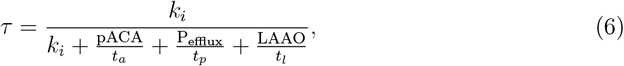

where *t*_*l*_, *t*_*a*_, and *t*_*p*_ are chemical-specific toxicity factors. Enzyme-induced toxicity *t*_*l*_ scales the key metabolite depletion rate driven by the enzyme LAAO. Metabolite-induced toxicity *t*_*a*_ scales the impact of toxic intermediate p-ACA concentration. Finally, protein-induced toxicity *t*_*p*_ reflects the toxicity caused by efflux pump expression. The toxicity factor acts as a scaling coefficient on the pathway synthesis, degradation, and folding reaction rates.

Despite the complexity and size of the p-AS model, we observe that BayesOpt explores many of the 27 possible architectures and converges to a low value of the objective function (Figure 4B); this was also achieved at a reasonable computational cost (mean run time under two minutes). The best architecture selected in the sample run was a double upstream repression, single downstream activation loop controlled by p-AF (Figure 4B, inset), but there is no clear best architecture when the optimization is run many times. No architecture is optimal for more than 15% of test runs, demonstrating that there are combinations of architectures and parameter values that achieve a similar optimal loss. We also found that several architectures can display oscillatory solutions, which we chose to exclude from the search by applying a peak detection algorithm (*48*) and adding a large regularization term to the loss.

To investigate the robustness to chemical toxicity, we perturbed the metabolite-induced toxicity *t*_*a*_ and protein-induced toxicity *t*_*p*_ in Eq. (6). The optimal loss values were found to be comparable between perturbed and background systems (Figure 4C). Additionally, when projected onto a 2-dimensional space using principal component analysis, the distribution of background parameter values was similar to the distribution of perturbed solutions, indicating that the perturbation did not significantly affect the optimal parameters selected (Figure 4D) (*49*). The p-AS pathway lies at the far end of what is currently possible to build experimentally and thus illustrates the broad applicability of BayesOpt to realistic design tasks in metabolic engineering.

## 3 Discussion

Progress in synthetic biology allows the construction of circuits on increased complexity and acting across various levels of biological organization. However, large design spaces and multiple scales of biological organization can become substantial challenges for the rapid design of functional systems. In this paper we presented the use of Bayesian optimization for the design of biological circuits. The method can rapidly find circuit architectures and parameters that optimize a performance objective that captures the target circuit functionality.

The method is particularly well suited for cases in which the multiple scales prevent efficient simulation of ODE models. Gene circuits designed to control metabolic pathways are an excellent example of such multiscale systems, as they combine fast metabolic timescales with the much slower dynamics of gene expression. Moreover, the choice of regulators, control points, and control architectures adds multiple degrees of freedom that are infeasible to explore experimentally. Previously implemented metabolic control systems have been built primarily based on application-specific knowledge of pathway features (*27, 50*). We have shown that Bayesian optimization can aid the design of such systems prior to implementation and serve as tools for *in silico* screening of competing designs that may have similar performance but entail different cost of wetlab implementation. We showed the efficiency and scalability of the method in several real-world case studies from metabolic engineering. In particular, the p-aminostyrene pathway is more complex than systems typically implemented in literature so far, which suggests that the method is applicable across a range of real-world design tasks.

We anticipate several novel applications of this work to other problem areas where discovery or tuning of multiscale circuits has been previously infeasible. For instance, this method could be employed to fit temporal circuit dynamics to data or discern which of several discrete circuit mechanisms most closely matches observed behavior. As with other design strategies based on ODE models, a challenge of our approach is the significant domain knowledge required to construct models for a target pathway, both in terms of the enzyme kinetics and the downstream metabolic processes that affect pathway activity. Machine learning has already proved useful in a range of metabolic engineering tasks (*51*) and is gaining substantial interest in other areas of synthetic biology (*52, 53*). In this paper we have shown how such methods can also benefit dynamic pathway engineering by using optimization as a means to navigate the design space prior to system prototyping.

## 4 Methods

### 4.1 Bayesian optimization

We employed the Bayesian optimization routine implemented in the Python `HyperOpt` package (*37*). Bayesian optimization is commonly employed for hyperparameter tuning in deep neural networks. We employed Expected Improvement as an acquisition function and a tree-structured Parzen estimator (TPE) as a non-parametric statistical model for the loss landscape. We performed a grid search over the TPE hyperparameter *γ* which controls the balance between exploration and exploitation but found little impact on the algorithm performance; we thus used the default value of *γ* = 15 (Supplementary Figure S1).

Constraints on the continuous and discrete decision variables were incorporated directly into the `HyperOpt` search space. At each run of the Bayesian optimization routine, the initial guess for the continuous decision variables were sampled from uniform distributions, with upper and lower bounds were taken from literature (*10, 34, 44*). Architectures were chosen uniformly from the set of architectures without positive feedback loops.

### 4.2 Model pathways

We considered four exemplar pathways modelled via ordinary differential equations (ODEs): the toy system in Figure 1C, the glucaric acid pathway in Figure 2A, the fatty acid pathway in Figure 3A, and the p-aminostyrene pathway in Figure 4A. Table 1 contains a summary of the four considered models. In all cases, pathway models include ODEs for both metabolites and pathway enzymes. In each case, we define the various control architectures and incorporate them as discrete decision variables in the optimization problem, i.e. *p*_*d*_ in Eq. (1); the continuous decision variables, i.e. *p*_*c*_ in Eq. (1), appear in the expression rates of the pathway enzymes. For the toy model and the glucaric acid pathway, enzyme expression was parameterized using a lumped Hill equation model to describe the interaction between a regulatory metabolite and a transcription factor. For the fatty acid and p-aminostryrene pathways, expression rates were parameterized with bespoke nonlinear functions describing specific biochemical processes. The discrete control architectures were defined in two different ways. For the toy, glucaric acid, and p-aminostyrene models, the architectures were defined using a binary matrix to encode the mode of transcriptional control. For the fatty acid model we instead defined each architecture as a categorical choice and switched between model functions correspondingly. We note that the p-aminostyrene pathway also contains ODEs for mRNA abundance and folded/unfolded proteins. All models and their parameters are described in the Supplementary Information.

**Table 1:**
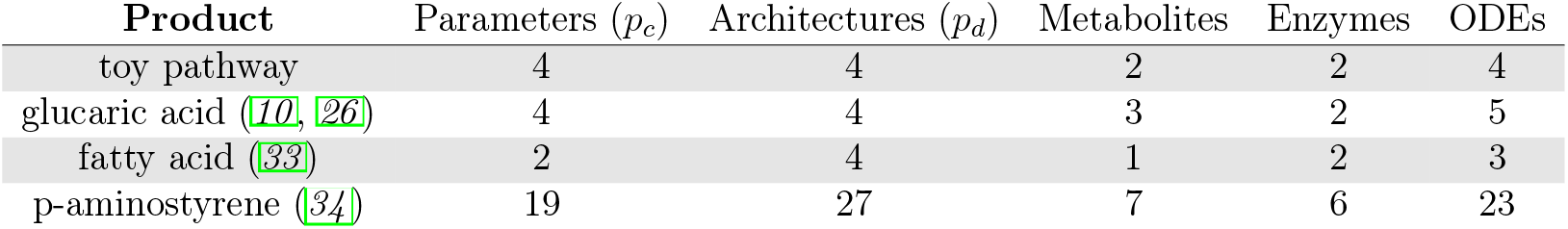
Summary of pathway models studied in this paper. The ODEs in the paminostyrene pathway also include mRNA and folding dynamics.

The ODE models were solved with `scikit-odes`, a Python wrapper for the sundials suite of solvers (*54*). In all cases, the initial concentrations of heterologous pathway enzymes were assumed to be zero. Initial concentrations for native metabolites were determined by first solving a model without the heterologous enzymes up to steady state. Simulation times and initial conditions are detailed in the Supplementary Information for each model.

### 4.3 Loss function

In all cases the loss function *J* in Eq. (3) was instanced to each specific pathway. Generally, the loss is defined as a linear combination of costs and benefits of pathway activity so as to balance opposing design goals commonly found in applications. Since both components of the loss function have different magnitudes, for each model we first swept the weights *α*_1_ and *α*_2_ across many model simulations, and chose values that led to similar values for both components; this prevents the optimizer to bias the search towards low loss values caused by the scaling effects. For the fatty acid model in Figure 3 we also optimized the circuit rise-time and overshoot % defined in Eq. (5). Details on all objective functions can found in the Supplementary Information.

## Code availability

Python code for this paper is available on Zenodo at https://doi.org/10.5281/zenodo.7926205

## Supporting information

Details of model construction for toy, glucaric acid, fatty acid, and p-aminostyrene models, including parameter values and model equations; Additional supporting figures related to hyperparameter tuning.

## Author information

### Author contributions

CM built the optimization pipeline, ran simulations, and produced figures. CM and DAO analyzed results. DAO and OMA supervised the research.

