## Supplementary Information for "Bayesian optimization for design of multiscale biological circuits"

#### S1 Overview of Pathways Under Study

We chose four experimentally implemented pathways with existing peer-reviewed models from literature to demonstrate different potential challenges and applications for the method. Details of the differential equation implementation are given below. A summary of relevant statistics for each method is shown in Table 1. Optimizable parameters for each model are highlighted in red.

#### S2 Toy Pathway

The first pathway under consideration is a toy pathway with two pathway intermediates and enzymes under transcriptional control (*10*). The substrate represses a metabolite-responsive transcription factor (MRTF) which can act as an activator or repressor for the transcription of enzymes 1 and 2. (see Figure 1A). The influx from native precursors and draw from pathway metabolism are modeled as constant rates. While there is no direct biological correlate for this pathway, it exemplifies the most basic engineered pathway possible and thus is useful primarily as a proof-of-concept for the BayesOpt method.

The mass balance equations for the toy branched pathway are

$$\begin{aligned}
\frac{\mathrm{d}x_0}{\mathrm{d}t} &= V_{\mathrm{in}} - e_0\frac{k_{\mathrm{cat}}x_0}{k_{\mathrm{m}}+x_0} - e_1\frac{k_{\mathrm{cat}}x_0}{k_{\mathrm{m}}+x_0} - \lambda x_0,\\
\frac{\mathrm{d}x_1}{\mathrm{d}t} &= e_1\frac{k_{\mathrm{cat}}x_0}{k_{\mathrm{m}}+x_0} - e_2\frac{k_{\mathrm{cat}}x_1}{k_{\mathrm{m}}+x_1} - \lambda x_1,\\
\frac{\mathrm{d}e_1}{\mathrm{d}t} &= u(x_1, k_1, \theta_1, n) - \lambda e_1,\\
\frac{\mathrm{d}e_2}{\mathrm{d}t} &= u(x_1, k_2, \theta_2, n) - \lambda e_2,
\end{aligned} \tag{S1}$$

where $x_0$ and $x_1$ are the metabolite concentrations and $e_1$ and $e_2$ are the enzyme concentrations. The input flux $V_{\mathrm{in}}$ is the flux from native precursors. The constant $\lambda$ is the culture growth rate and is fixed at $1.93 \cdot 10^-4s^-1$, which corresponds to a 26 minute doubling time in *E. coli*. Each enzyme follows standard Michaelis-Menten kinetics has two fixed kinetic parameters, $k_{\mathrm{cat}}$ and $k_{\mathrm{m}}$. For simplicity, all enzymes are assumed to have the same kinetic parameter values. The function $u(x, k_i, \theta_i, n)$ describes the expression rates of each enzyme, which can be modeled by one of three functions dependent on genetic control architecture (se1e Equation S2). The dose-response curves are encoded by the optimizable parameters $k_i$ and $\theta_i$ and the fixed Hill coefficient $n = 2$. The form of $u(x, k, \theta)$ is determined by the architecture value and modeled using sigmoid functions which lump together multiple molecular processes, including metabolite-TF and TF-DNA binding. These lumped models are simpler to parameterize using limited kinetic data. Additionally, dynamic pathways often express the TF constitutively, which reduces the effect of TF expression on performance ([24]). We consider three architecture forms: activation, repression, and open loop control:

$$u(x_i, k_i, \theta_i) = \begin{cases} k_i & \text{(no control)},\\ \dfrac{k_i x^2}{{\theta_i}^2 + x^2} & \text{(activation)},\\ \dfrac{k_i {\theta_i}^2}{{\theta_i}^2 + x^2} & \text{(repression)}.\end{cases} \tag{S2}$$

As a result, the toy pathway has four continuous parameters under study: $k_1$, $\theta_1$, $k_2$, $\theta_2$, two associated with each enzyme. The four optimizable parameters are constrained on the following ranges:

$$10^{-3} \leq \theta_1, \theta_2 \leq 10, \tag{S3}$$

$$10^{-7} \leq k_1, k_2 \leq 10^{-3}. \tag{S4}$$

The possible architectures are limited to three architectures which include only negative feedback loops, in addition to an open-loop control with no dynamic feedback loops (see Figure 1A). This architecture limitation was made to remove the possibility of undesirable multistability or oscillatory behavior. All parameter values are given in Table S1.

Supplementary Table S1: Parameter values of the toy model in Equation S1. All model values come from (*10*).

| Parameter | Value | Units |
|---|---|---|
| $k_{\text{cat}}$ | 12 | $\mu$M/s |
| $k_{\text{m}}$ | 10 | $\mu$M |
| $V_{\text{in}}$ | 1 | $\mu$M/s |
| $e_0$ | 0.0467 | $\mu$M |
| $\lambda$ | 1.93E-4 | 1/s |
| $\alpha_1$ | 1E-5 | N/A |
| $\alpha_2$ | 1E-2 | N/A |

The $\alpha_1$ and $\alpha_2$ values given in Table S1 were chosen to scale the two objectives to be similar in magnitude and the overall loss to be between 0 and 1. The loss equation for the objective function is

$$J = \underbrace{\alpha_1 \int_0^T \left| V_{\text{in}} - e_2(t) \frac{k_{\text{cat}} x_1(t)}{k_{\text{m}} + x_1(t)} \right| \mathrm{d}t}_{\text{production loss}} + \underbrace{\alpha_2 \int_0^T \left( u(x_1(t), k_1, \theta_1) + u(x_1(t), k_2, \theta_2) \right) \mathrm{d}t}_{\text{pathway cost}} . \tag{S5}$$

We ran each simulation of the model for a total time of $5 \cdot 10^4$ seconds to ensure the final values would be at steady state. All metabolites and enzymes had an initial concentration of 0$\mu$M except for $x_0$, which had an initial concentration of 2290$\mu$M (*10*). A summary of the model details is given in Table S2.

Supplementary Table S2: Toy model summary. The names for the decision variables and pathway metabolites and enzymes are included, as are the initial condition values.

| **Pathway Product** | Toy Product |
|---|---|
| Decision Variables | $k_1, k_2, \theta_1, \theta_2$ |
| Architectures | Open Loop, Upstream Repression, Downstream Activation, Dual Control |
| Pathway Metabolites | $X_0, X_1$ |
| Pathway Enzymes | $E_0$ (constant), $E_1, E_2$ |
| Integration Time | $5 \cdot 10^4$s |
| **Initial Conditions** | $x_0(0) = 2290\mu$M<br>$x_1(0) = 0\mu$M<br>$e_1(0) = 0\mu$M<br>$e_2(0) = 0\mu$M |

### S3 Glucaric Acid Production

The second pathway describes the synthesis of glucaric acid, a key precursor for a number of applications (36). This pathway has been implemented using dynamic control architectures which led to a 2.5-fold increase in product titer over static metabolic engineering methods (26). Glucose is converted to glucose-6-phosphate, which can either be drawn into native metabolism and converted, reversibly, into fructose-6-phosphate, or into the engineered pathway (see Figure 2A). The pathway requires three foreign enzymes: inositoal-3-phosphate synthetase, or Ino1, from *Saccharomyces cerevisiae*, myoinositol oxidase (MIOX), from *Mus musculus*, and uronate dehydrogenase (Udh) from *Pseudomonas syringae*. In addition to being exported from the cell and acting allosterically on MIOX, myoinositol can sequester the transcription factor IpsA, a dual transcriptional regulator from *Corynebacterium glutamicum*. IpsA can then act on the transcription of Ino1 or MIOX. The glucaric acid pathway provides a more complex, real-world application that builds on the toy model and includes reversible and allosteric reactions.

The glucaric acid model was chosen for its increased complexity. While maintaining the same number of possible architectures as the toy model, this model incorporates allosteric control and reversible reactions, two more complex biological interactions. We specify the model equations and describe a sample optimization. We then experimented with perturbing the growth conditions and kinetic parameters to investigate model robustness.

The glucaric acid pathway is based on the one designed in (*10*). The mass balance equations for the glucaric acid production pathway (see Figure 2A) are

$$
\begin{aligned}
\frac{\mathrm{dg6p}}{\mathrm{d}t} &= V_{\mathrm{in}} - \mathrm{zwf}\frac{k_{\mathrm{cat,\ zwf}}\mathrm{g6p}}{k_{\mathrm{m,\ zwf}} + \mathrm{g6p}} - \mathrm{pgi}\frac{k_{\mathrm{cat,\ pgi}}(\mathrm{g6p} - (\mathrm{f6p}/k_{\mathrm{eq}}))}{\mathrm{g6p} + k_{\mathrm{m,\ pgi,\ g6p}}(1 + (\mathrm{f6p}/k_{\mathrm{m,\ pgi,\ f6p}}))} - \lambda \cdot \mathrm{g6p},\\
\frac{\mathrm{df6p}}{\mathrm{d}t} &= \mathrm{pgi}\frac{k_{\mathrm{cat,\ pgi}}(\mathrm{g6p} - (\mathrm{f6p}/k_{\mathrm{eq}}))}{\mathrm{g6p} + k_{\mathrm{m,\ pgi,\ g6p}}(1 + (\mathrm{f6p}/k_{\mathrm{m,\ pgi,\ f6p}}))} + 0.5\mathrm{zwf}\frac{k_{\mathrm{cat,\ zwf}}\mathrm{g6p}}{k_{\mathrm{m,\ zwf}}} - \frac{k_{\mathrm{cat,\ pfk}}\mathrm{f6p}^3}{k_{\mathrm{m,\ pfk}}^3 + \mathrm{f6p}^3} - \lambda \cdot \mathrm{f6p},\\
\frac{\mathrm{dMI}}{\mathrm{d}t} &= \mathrm{ino1}\frac{k_{\mathrm{cat,\ ino1}}\mathrm{g6p}}{k_{\mathrm{m,\ ino1}}} - \frac{V_{\mathrm{m,\ t,\ MI}}\mathrm{MI}}{k_{\mathrm{m,\ t,\ MI}} + \mathrm{MI}} - \mathrm{MIOX}\frac{k_{\mathrm{cat,\ eff}}\mathrm{MI}}{k_{\mathrm{m,\ MIOX}} + \mathrm{MI}} - \lambda \cdot \mathrm{MI},\\
\frac{\mathrm{dino1}}{\mathrm{d}t} &= u(\mathrm{MI}, k_{\mathrm{ino1}}, \theta_{\mathrm{ino1}}) - \lambda \cdot \mathrm{ino1},\\
\frac{\mathrm{dMIOX}}{\mathrm{d}t} &= u(\mathrm{MI}, k_{\mathrm{MIOX}}, \theta_{\mathrm{MIOX}}) - \lambda \cdot \mathrm{MIOX},
\end{aligned}
\tag{S6}
$$

where Ino1 and MIOX are the enzymes in the pathway and g6p, f6p, and MI are the substrates. The enzyme parameters $k_{\mathrm{cat}}, k_{\mathrm{eq}}, k_{\mathrm{m,\ x}}$ and $k_{\mathrm{m,\ y}}$ are all fixed kinetic parameters specific to each enzyme. The effective substrate activation constant $k_{\mathrm{cat,\ eff}}$ has two additional activation kinetic parameters $k_{\mathrm{a}}$ and $a$ which must be specified:

$$
k_{\mathrm{cat,\ eff}} = k_{\mathrm{cat,\ MIOX}}\frac{1 + a_{\mathrm{MIOX}}\mathrm{MI}}{k_{\mathrm{a,\ MIOX,\ MI}} + \mathrm{MI}}, \tag{S7}
$$

The function $u(x, k, \theta)$ describes the genetic control topology at the enzyme's promoter. There are three options for this functional form: activation, repression, and no control as in the toy model (see Equation S2).

Similarly to the toy pathway, there are four continuous parameters: $k_{\mathrm{ino1}}$, $\theta_{\mathrm{ino1}}$, $k_{\mathrm{MIOX}}$, $\theta_{\mathrm{MIOX}}$. The four decision variables are constrained on the following ranges:

$$
1 \cdot 10^{-7} \leq \theta_{\mathrm{ino1}}, \theta_{\mathrm{MIOX}} \leq 10, \tag{S8}
$$

$$
1 \cdot 10^{-7} \leq k_{\mathrm{ino1}}, k_{\mathrm{MIOX}} \leq 5. \tag{S9}
$$

The possible architectures are limited to those only containing negative feedback loops. Similarly to the toy model, we term these three architectures upstream repression, downstream activation, and dual control based on their mode of action (see Figure 2A). All kinetic parameters are given in Table S3.

Supplementary Table S3: Glucaric acid model kinetic parameters.

| **Parameter** | **Value** | **Units** | **Parameter** | **Value** | **Units** |
|---|---|---|---|---|---|
| $\lambda$ | 2.77E-5 | 1/s | $n_{\text{Pfk}}$ | 3 | N/A |
| $V_{\text{in}}$ | 0.1656 | mM/s | $k_{\text{cat, ino1}}$ | 0.2616 | mM/s |
| $k_{\text{cat, pgi}}$ | 0.8751 | mM/s | $k_{\text{m, ino1, g6p}}$ | 1.18 | mM |
| $k_{\text{eq, pgi}}$ | 0.3 | mM | $V_{\text{m, t, MI}}$ | 0.045 | mM/s |
| $k_{\text{m, pgi, g6p}}$ | 0.28 | mM | $k_{\text{m, t, MI}}$ | 15 | mM |
| $k_{\text{m, pgi, f6p}}$ | 0.147 | mM | $k_{\text{cat, MIOX}}$ | 0.2201 | mM/s |
| $k_{\text{cat, zwf}}$ | 0.0853 | mM/s | $k_{\text{m, MIOX}}$ | 24.7 | mM |
| $k_{\text{m, zwf, g6p}}$ | 0.1 | mM | $a_{\text{MIOX}}$ | 5.422 | N/A |
| $k_{\text{cat, pfk}}$ | 2.615 | mM/s | $k_{\text{a, MIOX, MI}}$ | 20 | mM |
| $k_{\text{m, pfk, f6p}}$ | 0.16 | mM | $\alpha_1$ | 10E-5 | N/A |
| | | | $\alpha_2$ | 10E-3 | N/A |

Supplementary Table S4: Glucaric acid model summary. The names for the decision variables and pathway metabolites and enzymes are included, as are the initial condition values. All model values come from (*10*).

| **Pathway Product** | Glucaric Acid |
|---|---|
| Decision Variables | $k_{\text{ino1}}, k_{\text{MIOX}}, \theta_{\text{ino1}}, \theta_{\text{MIOX}}$ |
| Architectures | Open Loop, Upstream Repression, Downstream Activation, Dual Control |
| Pathway Metabolites | g6p, f6p, MI |
| Pathway Enzymes | Pgi (constant), Zwf (constant), Pfk (constant), Ino1, MIOX |
| Integration Time | $5 \cdot 10^5$s |
| **Initial Conditions** | f6p(0) $=$ 0.281mM<br>g6p(0) $=$ 0.0605mM<br>MI(0) $=$ 0mM<br>Ino1(0) $=$ 0mM<br>MIOX(0) $=$ 0mM |

The $\alpha_1$ and $\alpha_2$ values in the objective function were chosen to scale the two objectives to be similar in magnitude and the overall loss to be between 0 and 1 (see Table S3). The loss equation for the objective function is

$$J = \alpha_1 \underbrace{\int_0^T \left| V_{\text{in}} - \text{MIOX}(t) \frac{k_{\text{cat, MIOX}} \text{MI}(t)}{k_{\text{m, MIOX}} + \text{MI}(t)} \right| \mathrm{d}t}_{\text{production loss}} \tag{S10}$$

$$+ \alpha_2 \underbrace{\int_0^T \left( u(\text{MI}(t), k_{\text{ino1}}, \theta_{\text{ino1}}) + u(\text{MI}(t), k_{\text{MIOX}}, \theta_{\text{MIOX}}) \right) \mathrm{d}t}_{\text{pathway cost}} . \tag{S11}$$

A summary of the model details is given in Table S4. We ran each simulation of the model to a final time of $5 \cdot 10^5$s to ensure the final values would be at steady state. All metabolites and enzymes had an initial concentration of $0\mu$M except for g6p and f6p, which had initial concentrations of 0.281mM and 0.0605mM, respectively. These values were obtained by running the model with the engineered pathway removed and taking the steady-state values of the unmodified pathway metabolites as a baseline.

### S4 Fatty Acid Synthesis

Fatty acids are one of the four macromolecule types necessary for life. In addition to being used to form cell membranes, they are important sources of energy. The engineered fatty acid biosynthetic pathway shown in Figure 3A was created by expressing a thioesterase under transcriptional control (*33*). The architectures implemented in this pathway include both negative feedback loops built with a metabolite-responsive transcription factor as well as a genetic feedback loop where a repressor is produced on the same promoter as the enzyme.

The fatty acid model has multiple different architectures which model different molecular components of the pathway. Due to this model complexity, we consider each architecture in detail. The mass balance equations for the open loop architecture are

$$\begin{aligned} \frac{\text{dFFA}}{\mathrm{d}t} &= k_{\text{tesA}} \cdot \text{tesA} - \lambda \cdot \text{FFA}, \\ \frac{\text{dtesA}}{\mathrm{d}t} &= r_{\text{lac}} - \lambda \cdot \text{tesA}, \end{aligned} \tag{S12}$$

where FFA is the concentration of free fatty acids and tesA is the concentration of thioesterase enzyme (tesA). The cellular growth rate $\lambda$ is set for all architectures at $3.85 \cdot 10^{-4}$ mM/s, which is calculated from an *E. coli* doubling time of 26 minutes. The parameter $k_{\mathrm{tesA}}$ is the kinetic parameter of the thioesterase reaction. There is one free optimisable parameter in this model, the tesA promoter constant $r_{\mathrm{lac}}$.

The negative gene loop architecture expresses a transcriptional repressor on the same promoter as tesA, creating a negative feedback loop on enzyme production. The mass balance equations for this architecture are

$$\begin{aligned} \frac{\mathrm{dFFA}}{\mathrm{d}t} &= k_{\mathrm{tesA}} \cdot \mathrm{tesA} - \lambda \cdot \mathrm{FFA}, \\ \frac{\mathrm{dtesA}}{\mathrm{d}t} &= \frac{r_{\mathrm{tl}}\mathrm{tetR}^2}{k_{\mathrm{d,\ tetR}}^2 + \mathrm{tetR}^2} - \lambda \cdot \mathrm{tesA}, \\ \frac{\mathrm{dtetR}}{\mathrm{d}t} &= \frac{r_{\mathrm{tl,\ tetR}}\mathrm{tetR}^2}{k_{\mathrm{d,\ tetR}}^2 + \mathrm{tetR}^2} - \lambda \cdot \mathrm{tetR}. \end{aligned} \tag{S13}$$

The variable tetR is the concentration of the repressor expressed on the same gene as tesA. While the other models allow multiple modes of transcriptional control (activation, repression, no control), this model only allows for activation due to the type of transcription factor used ([33]). The relative expression strengths $r_{\mathrm{tl}}$ and $r_{\mathrm{tl,\ tetR}}$ are free parameters.

The negative metabolic loop architecture includes a transcription factor which acts on the tesA promoter. The mass balance equations for this architecture are

$$\begin{aligned} \frac{\mathrm{dFFA}}{\mathrm{d}t} &= k_{\mathrm{tesA}} \cdot \mathrm{tesA} - \lambda \cdot \mathrm{FFA}, \\ \frac{\mathrm{dtesA}}{\mathrm{d}t} &= \frac{r_{\mathrm{fl'}}\mathrm{tetR}^2}{k_{\mathrm{i}}^2} - \lambda \cdot \mathrm{tesA}, \end{aligned} \tag{S14}$$

where the free parameters of this model are $k_i$ and $r_{\mathrm{fl}'}$, which control the shape of the transcription factor dose-response curve. Unlike the other free model parameters, $k_i$ varies on the range 0 to 0.12.

Finally, the layered negative feedback loop uses tetR as an intermediate repressor. The transcription factor is repressed by the product and in turn represses the repressor TetR, which can repress the expression of the enzyme tesA. This negative feedback loop is described by the following equations:

$$\begin{aligned}\frac{\mathrm{dFFA}}{\mathrm{d}t} &= k_{\mathrm{tesA}} \cdot \mathrm{tesA} - \lambda \cdot \mathrm{FFA},\\ \frac{\mathrm{dtesA}}{\mathrm{d}t} &= \frac{r_{\mathrm{tl}}\mathrm{tetR}^2}{k^2_{\mathrm{d,\ tetR}}} - \lambda \cdot \mathrm{tesA},\\ \frac{\mathrm{dtetR}}{\mathrm{d}t} &= \frac{r_{\mathrm{ar2}} k^2_{\mathrm{ar2}}}{(1+\frac{\mathrm{FFA}}{k_{\mathrm{d,\ FadR,\ FFA}}})^2} - \lambda \cdot \mathrm{tetR}.\end{aligned} \tag{S15}$$

The repressor TetR is produced on the AR2 promoter, which has its own strength parameter $r_{\mathrm{ar2}}$. The free parameters on this model are $r_{\mathrm{ar2}}$ and $r_{\mathrm{tl}}$.

Unless otherwise stated, all continuous model parameters are restricted to the range from $10 \cdot 10^{-11}$ to $10 \cdot 10^{-8}$. We ran each simulation of the model to $5 \cdot 10^4$s to ensure steady state values had been reached. All pathway components started with a concentration of 0mM. Two objective functions were implemented for this pathway: a production-burden function and a speed-accuracy function. The loss equation for the production-burden objective function is the sum of the production loss and pathway cost:

$$J = \alpha_1 J_{\mathrm{prod}} + \alpha_2 J_{\mathrm{cost}}, \tag{S16}$$

where $J_{\mathrm{prod}}$ is the production loss. The $\alpha_1$ and $\alpha_2$ values in the objective function were chosen to scale the two objectives to be similar in magnitude and the overall loss to be between 0 and 1 (see Table S3). The production loss is defined as

$$J_{\mathrm{prod}} = \frac{1}{\int_0^T |\mathrm{tesA}(t) k_{\mathrm{tesA}}| \ \mathrm{d}t}. \tag{S17}$$

The pathway cost $J_{\text{cost}}$ varies for each architecture:

$$J_{\text{cost}} = \begin{cases} \int_0^T (r_{\text{lac}})\,\mathrm{d}t & \text{(open loop)}, \\ \int_0^T (\frac{r_{\text{tl}}\text{tetR}^2}{k^2_{\text{d, tetR}} + \text{tetR}^2} + \frac{r_{\text{tl, tetR}}\text{tetR}^2}{k^2_{\text{d, tetR}} + \text{tetR}^2})\,\mathrm{d}t & \text{(negative gene loop)}, \\ \int_0^T (\frac{r_{\text{fl}}\text{tetR}^2}{k_{\text{i}}^2})\,\mathrm{d}t & \text{(negative metabolic loop)}, \\ \int_0^T (\frac{r_{\text{tl}}\text{tetR}^2}{k^2_{\text{d, tetR}}} + \frac{r_{\text{ar2}}k^2_{\text{ar2}}}{(1 + \frac{\text{FFA}}{k_{\text{d, FadR, FFA}}})^2})\,\mathrm{d}t & \text{(negative gene loop)}. \end{cases} \quad \text{(S18)}$$

The second speed-accuracy objective function has two terms, percent overshoot and normalized rise-time:

$$J = \alpha J_{\text{rt}} + J_{\text{os}}. \quad \text{(S19)}$$

The percent overshoot $J_{\text{os}}$ is the percent difference between the maximum FFA concentration and the steady-state FFA concentration at the end of the integration time. For example, a 10% overshoot would correspond to a maximum FFA concentration 10% over the steady-state concentration. The rise-time is the first time point where the FFA concentration reaches 50% of the steady-state value, normalized by the total integration time. As a result, a 20% rise-time corresponds to the FFA concentration reaching 50% of the steady-state value one-fifth of the way through the total integration time. Table S6 summarizes the model details.

### S5 P-Aminostyrene Synthesis

The final pathway we consider is the synthesis of p-aminostyrene (p-AS) in *E. coli* (*34*). p-AS is an industrially relevant vinyl aromatic monomer with applications in photonics and biomedicine (*45*). However, the cytotoxic intermediate p-amminocinnamic acid (p-ACA) makes p-AS challenging to produce in microbes. Furthermore, another intermediate, p-aminophenylalanine (p-AF), leaks from cells (*57*). L-Amino Acid Oxidase, one of the enzymes in the pathway (see Figure 4A) depletes key aromatic amino acid metabolites and creates toxic hydrogen peroxide as a byproduct. Finally, overexpression of the efflux pump

Supplementary Table S5: Fatty acid model kinetic parameters

| Architecture | Parameter | Definition | Value | Units |
|---|---|---|---|---|
| All | $\lambda$ | growth rate | $3.85 \cdot 10^{-4}$ | mM/s |
| OL | $k_{\text{tesA}}$ | tesA catalysis constant | 100 | mM/s |
| NGL | $k_{\text{tesA}}$ | tesA catalysis constant | 105.25 | mM/s |
| NGL | $k_{\text{d, tetR}}$ | tetR dissociation constant | $3.0 \cdot 10^{-8}$ | mM/s |
| NML | $k_{\text{tesA}}$ | tesA catalysis constant | 77.75 | mM/s |
| LNML | $k_{\text{tesA}}$ | tesA catalysis constant | 230.9 | mM/s |
| LNML | $k_{\text{d, tetR}}$ | tetR dissociation constant | $3.85 \cdot 10^{-8}$ | mM/s |
| LNML | $k_{\text{d, FadR, FFA}}$ | tetR and FFA dissociation constant | 0.001 | mM/s |
| LNML | $k_{ar2}$ | tetR promoter constant | 138.50 | mM/s |
| all | $\alpha_1$ | production loss scaling constant | 10E-4 | mM/s |
| all | $\alpha_2$ | pathway cost scaling constant | 10E-3 | mM/s |

Supplementary Table S6: Fatty acid model summary. Some metabolites and enzymes (tetR) are only modeled for relevant architectures like negative metabolic loop.

| **Pathway Product** | Fatty Acid |
|---|---|
| Decision Variables | $r_{\text{lac}}, r_{\text{bad}}, r_{\text{tl}}, r_{\text{tl, tetR}}, r_{\text{fl'}}, r_{\text{ar2}}$ |
| Architectures | Open Loop, Negative Gene Loop, Negative Metabolic Loop, Layered Negative Metabolic Loop |
| Pathway Metabolites | FFA, tetR (repressor) |
| Pathway Enzymes | tesA |
| Integration Time | $5 \cdot 10^4$s |
| **Initial Conditions** | 0mM for all metabolites and enzymes |

used to remove p-ACA from the cell also causes cytotoxicity. In addition to these challenges, the pathway has two possible loci of genetic control. Both p-AF and p-ACA can act as ligands for aptazyme-regulated expression device (aRED) transcription factors. These aRED transcription factors can then act as dual activator-repressors on any of the three promoters involved in the pathway. The first three enzymes in the pathway, papA, papB, and papC, are all expressed on the papABC operon, which produces a single mRNA transcript. Operons are common in bacteria, so including one in our pathway models demonstrates an important application of this method. The p-aminostryene synthesis model is the largest and most complex pathway under study. It includes two metabolites which can act on transcription factors: p-ACA and p-AF. There are a large number of possible architectures possible for this model (27 excluding positive feedback loops). As a result, we do not explicitly name the architectures.

The P-aminostyrene pathway is based on one studied in (*34*). There are 7 pathway metabolites, 3 DNA promoter elements, 5 mRNA transcripts, 4 unfolded enzymes, 5 folded enzymes, and a folded and unfolded efflux pump protein described in the mass balance equations. We modified the Stevens model to simplify the multiple explicit equations describing aRED aptamer folding into a single sigmoid similar to those in Equation S2. The metabolite mass balance equations are

$$\begin{aligned}
\frac{\mathrm{dchorismate}}{\mathrm{d}t} &= V_{\mathrm{chorismate}}\tau - f(\mathrm{papA}, \mathrm{chorismate}) - \delta \cdot \mathrm{chorismate}, \\
\frac{\mathrm{dpA1}}{\mathrm{d}t} &= f(\mathrm{papA}, \mathrm{chorismate}) - f(\mathrm{papB}, \mathrm{pA1}) - \delta \cdot \mathrm{pA1}, \\
\frac{\mathrm{dpA2}}{\mathrm{d}t} &= f(\mathrm{papB}, \mathrm{pA1}) - f(\mathrm{papC}, \mathrm{pA2}) - \delta \cdot \mathrm{pA2}, \\
\frac{\mathrm{dpA3}}{\mathrm{d}t} &= f(\mathrm{papC}, \mathrm{pA2}) - f(\mathrm{deaminase}, \mathrm{pA3}) - \delta \cdot \mathrm{pA3}, \\
\frac{\mathrm{dpAF}}{\mathrm{d}t} &= f(\mathrm{deaminase}, \mathrm{pA3}) - f(\mathrm{LAAO}, \mathrm{pAF}) - \delta \cdot \mathrm{pAF} - \mathrm{L}, \\
\frac{\mathrm{dpACA}}{\mathrm{d}t} &= f(\mathrm{LAAO}, \mathrm{pAF}) - f(\mathrm{P}_{\mathrm{efflux}}, \mathrm{pACA}) - \delta \cdot \mathrm{pACA}, \\
\frac{\mathrm{dpAS}}{\mathrm{d}t} &= f(\mathrm{P}_{\mathrm{efflux}}, \mathrm{pACA}).
\end{aligned} \tag{S20}$$

Here, the constant L is the rate of loss of p-AF through the leaky cellular membrane, and $V_{\mathrm{chorismate}}$ is a constant parameter defined in Table S8. The constant $\tau$ is the toxicity factor which scales concentrations to account for intermediate metabolite toxicity. The constant $\delta$ is the dilution rate due to cellular growth. The function $f$ is a variation on the Michaelis-Menten equation:

$$f(e,x) = k_{\text{cat}} e \frac{\frac{x}{\text{N}_\text{A}\text{Vol}_\text{cell}}}{k_\text{m} + \frac{x}{\text{N}_\text{A}+\text{Vol}_\text{cell}}} \tau,$$

where: (S21)

$e \in \{\text{papA, papB, papC, deaminase, LAAO, P}_\text{efflux}\}$,

$x \in \{\text{chorismate, pA1, pA2, pA3, pAF, pACA}\}$.

The promoter and mRNA transcript concentrations are modeled separately here, unlike in the previous models. The mass-balance equations for these elements are

$$\begin{aligned}
\frac{\text{dPr}_1}{\text{d}t} &= \text{Pr}_1\beta - \delta \cdot \text{Pr}_1,\\
\frac{\text{dPr}_2}{\text{d}t} &= \text{Pr}_2\beta - \delta \cdot \text{Pr}_2,\\
\frac{\text{dPr}_3}{\text{d}t} &= \text{Pr}_3\beta - \delta \cdot \text{Pr}_3,\\
\frac{\text{dmRNA}_\text{papA}}{\text{d}t} &= \omega_1 u(\text{pAF}, k_{1,\text{pAF}}, \theta_{1,\text{pAF}}) + (1-\omega_1) u(\text{pACA}, k_{1,\text{pACA}}, \theta_{1,\text{pACA}})\\
&\quad - \delta \cdot \text{mRNA}_\text{papA} - \tau \cdot \mu \cdot \text{mRNA}_\text{papA},\\
\frac{\text{dmRNA}_\text{papB}}{\text{d}t} &= \omega_1 u(\text{pAF}, k_{2,\text{pAF}}, \theta_{1,\text{pAF}}) + (1-\omega_1) u(\text{pACA}, k_{2,\text{pACA}}, \theta_{1,\text{pACA}})\\
&\quad - \delta \cdot \text{mRNA}_\text{papB} - \tau \cdot \mu \cdot \text{mRNA}_\text{papB},\\
\frac{\text{dmRNA}_\text{papC}}{\text{d}t} &= \omega_1 u(\text{pAF}, k_{3,\text{pAF}}, \theta_{1,\text{pAF}}) + (1-\omega_1) u(\text{pACA}, k_{3,\text{pACA}}, \theta_{1,\text{pACA}})\\
&\quad - \delta \cdot \text{mRNA}_\text{papC} - \tau \cdot \mu \cdot \text{mRNA}_\text{papC},\\
\frac{\text{dmRNA}_\text{LAAO}}{\text{d}t} &= \omega_2 u(\text{pAF}, k_{4,\text{pAF}}, \theta_{2,\text{pAF}}) + (1-\omega_2) u(\text{pACA}, k_{4,\text{pACA}}, \theta_{2,\text{pACA}})\\
&\quad - \delta \cdot \text{mRNA}_\text{LAAO} - \tau \cdot \mu \cdot \text{mRNA}_\text{LAAO},\\
\frac{\text{dmRNA}_{\text{P}_\text{efflux}}}{\text{d}t} &= \omega_3 u(\text{pAF}, k_{5,\text{pAF}}, \theta_{3,\text{pAF}}) + (1-\omega_3) u(\text{pACA}, k_{5,\text{pACA}}, \theta_{3,\text{pACA}})\\
&\quad - \delta \cdot \text{mRNA}_{\text{P}_\text{efflux}} - \tau \cdot \mu \cdot \text{mRNA}_{\text{P}_\text{efflux}},
\end{aligned} \tag{S22}$$

where the function $u(x, k, \theta)$ is the sigmoidal control function defined in Equation S2. The binary parameters $w_i$ define which metabolite exerts control on each promoter:

$$\omega_1 = \begin{cases} 1 & \text{papABC controlled by p-AF,} \\ 0 & \text{papABC controlled by p-ACA,} \end{cases}$$
$$\omega_2 = \begin{cases} 1 & \text{LAAO controlled by p-AF,} \\ 0 & \text{LAAO controlled by p-ACA,} \end{cases} \tag{S23}$$
$$\omega_3 = \begin{cases} 1 & \text{P}_{\text{efflux}}\text{controlled by p-AF,} \\ 0 & \text{P}_{\text{efflux}}\text{controlled by p-ACS.} \end{cases}$$

To avoid circuits with multistable dynamics, we restrict the search to architectures without positive feedback loops. This means that we consider a total of $3^3 = 27$, i.e. those where:

- promoter papABC is either unregulated or repressed by one of the two intermediates (p-AF or p-ACA),
- promoter LAAO is unregulated, activated by p-AF, or repressed by p-ACA,
- promoter $P_{\text{efflux}}$ is either unregulated or activated by one of the two intermediates (p-AF or p-ACA).

The constant $\beta$ is the DNA duplication rate. The constant $\mu$ is the mRNA degradation rate constant, which is assumed to be constant for all mRNAs. PapA, PapB, and PapC are all expressed from the same promoter but their translation rates are variable, so their mRNAs are modeled separately. Finally, the protein folding process is modeled explicitly, with each of the 4 enzymes and the efflux pump having a folded and unfolded state. Additionally, the enzyme deaminase is expressed constitutively but its concentration rises to steady state and thus its dynamics are also modeled. The enzyme mass balance equations are

$$
\begin{aligned}
\frac{\mathrm{dpapA_{uf}}}{\mathrm{d}t} &= v(\mathrm{mRNA_{papA}}) - \eta \cdot \tau \cdot \mathrm{papA_{uf}} - \delta \cdot \mathrm{papA_{uf}} - \rho \cdot \tau \cdot \mathrm{papA_{uf}}, \\
\frac{\mathrm{dpapB_{uf}}}{\mathrm{d}t} &= v(\mathrm{mRNA_{papB}}) - \eta \cdot \tau \cdot \mathrm{papB_{uf}} - \delta \cdot \mathrm{papB_{uf}} - \rho \cdot \tau \cdot \mathrm{papB_{uf}}, \\
\frac{\mathrm{dpapC_{uf}}}{\mathrm{d}t} &= v(\mathrm{mRNA_{papC}}) - \eta \cdot \tau \cdot \mathrm{papC_{uf}} - \delta \cdot \mathrm{papC_{uf}} - \rho \cdot \tau \cdot \mathrm{papC_{uf}}, \\
\frac{\mathrm{dLAAO_{uf}}}{\mathrm{d}t} &= v(\mathrm{mRNA_{LAAO}}) - \eta \cdot \tau \cdot \mathrm{LAAO_{uf}} - \delta \cdot \mathrm{LAAO_{uf}} - \rho \cdot \tau \cdot \mathrm{LAAO_{uf}}, \\
\frac{\mathrm{P_{efflux,\ uf}}}{\mathrm{d}t} &= v(\mathrm{mRNA_{P_{efflux,\ uf}}}) - \eta \cdot \tau \cdot \mathrm{P_{efflux,\ uf}} - \delta \cdot \mathrm{P_{efflux,\ uf}} - \rho \cdot \tau \cdot \mathrm{P_{efflux,\ uf}}, \\
\frac{\mathrm{dpapA}}{\mathrm{d}t} &= \eta \cdot \tau \cdot \mathrm{papA_{uf}} - \delta \cdot \mathrm{papA} - \rho \cdot \tau \cdot \mathrm{papA}, \\
\frac{\mathrm{dpapB}}{\mathrm{d}t} &= \eta \cdot \tau \cdot \mathrm{papB_{uf}} - \delta \cdot \mathrm{papB} - \rho \cdot \tau \cdot \mathrm{papB}, \\
\frac{\mathrm{dpapC}}{\mathrm{d}t} &= \eta \cdot \tau \cdot \mathrm{papC_{uf}} - \delta \cdot \mathrm{papC} - \rho \cdot \tau \cdot \mathrm{papC}, \\
\frac{\mathrm{dLAAO}}{\mathrm{d}t} &= \eta \cdot \tau \cdot \mathrm{LAAO_{uf}} - \delta \cdot \mathrm{LAAO} - \rho \cdot \tau \cdot \mathrm{LAAO}, \\
\frac{\mathrm{dP_{efflux}}}{\mathrm{d}t} &= \eta \cdot \tau \cdot \mathrm{P_{efflux,\ uf}} - \delta \cdot \mathrm{P_{efflux}} - \rho \cdot \tau \cdot \mathrm{P_{efflux}}, \\
\frac{\mathrm{ddeaminase}}{\mathrm{d}t} &= V_{\mathrm{deaminase}} - \delta \cdot \mathrm{deaminase}.
\end{aligned}
\tag{S24}
$$

Here, the constant $\rho$ is the protein degradation rate and $\eta$ is the protein folding rate. The function $v$ is the translation rate equation:

$$
v(m) = \frac{m}{\mathrm{T_{init}} + \frac{\mathrm{L}_m}{\mathrm{R}}}, \tag{S25}
$$

where $\mathrm{T_{init}}$ is the transcription initiation rate, $\mathrm{L}_m$ is the length of the mRNA, and R is the ribosome elongation rate. The objective function for the p-AS model is

$$
\begin{aligned}
J = \alpha_1 & \int_0^T |V_{\text{chorismate}}\tau - f(\mathrm{P}_{\text{efflux}}, \mathrm{pACA})| \, \mathrm{d}t \\
& + \alpha_2 \int_0^T [(\omega_1 u(\mathrm{pAF}, k_{1,\ \mathrm{pAF}}, \theta_{1,\ \mathrm{pAF}}) + (1-\omega_1)u(\mathrm{pACA}, k_{1,\ \mathrm{pACA}}, \theta_{1,\ \mathrm{pACA}}) \\
& + (\omega_1 u(\mathrm{pAF}, k_{2,\ \mathrm{pAF}}, \theta_{1,\ \mathrm{pAF}}) + (1-\omega_1)u(\mathrm{pACA}, k_{2,pACA}, 1,\ \mathrm{pACA}) \\
& + (\omega_1 u(\mathrm{pAF}, k_{3,\ \mathrm{pAF}}, \theta_{1,\ \mathrm{pAF}}) + (1-\omega_1)u(\mathrm{pACA}, k_{3,\ \mathrm{pACA}}, \theta_{1,\ \mathrm{pACA}}) \\
& + (\omega_2 u(\mathrm{pAF}, k_{4,\ \mathrm{pAF}}, \theta_{2,pAF}) + (1-\omega_2)u(\mathrm{pACA}, k_{4,\ \mathrm{pACA}}, \theta_{2,\ \mathrm{pACA}}) \\
& + (\omega_3 u(\mathrm{pAF}, k_{5,\ \mathrm{pAF}}, \theta_{3,\ \mathrm{pAF}}) + (1-\omega_3)u(\mathrm{pACA}, k_{5,\ \mathrm{pACA}}, \theta_{3,\ \mathrm{pACA}})]\mathrm{d}t,
\end{aligned} \tag{S26}
$$

where pathway cost is the sum of all heterologous enzyme transcription rates across all loci and ligands, and the rate $f(\mathrm{P}_{\text{efflux}}, \mathrm{pACA})$ is defined in Equation S21. The constant $V_{\text{chorismate}}$ as well as any other fixed parameter values in the model are defined in Table S8. Table S7 summarizes the model details.

Supplementary Table S7: P-Aminostryene model summary. The initial conditions for all model components are set to 0mM. Architectures are unnamed and thus not listed here.

| **Pathway Product** | P-Aminostyrene |
|---|---|
| Decision Variables | $\theta_{1,\ \mathrm{pAF}}$, $\theta_{1,\ \mathrm{pACA}}$, $\theta_{2,\ \mathrm{pAF}}$, $\theta_{2,\ \mathrm{pACA}}$, $\theta_{3,\ \mathrm{pAF}}$, $\theta_{3,\ \mathrm{pACA}}$, $k_{1,\ \mathrm{pAF}}$, $k_{1,\ \mathrm{pACA}}$, $k_{2,\ \mathrm{pAF}}$, $k_{2,\ \mathrm{pACA}}$, $k_{3,\ \mathrm{pAF}}$, $k_{3,\ \mathrm{pACA}}$, $k_{4,\ \mathrm{pAF}}$, $k_{4,\ \mathrm{pACA}}$, $k_{5,\ \mathrm{pAF}}$, $k_{5,\ \mathrm{pACA}}$ |
| Pathway Metabolites | Chorismate, PA1, PA2, PA3, pAF, pACA, PAS |
| Pathway Enzymes | papA, papB, papC, LAAO, Efflux pump, Deaminase |
| Integration Time | $1.73 \cdot 10^5$s |
| **Initial Conditions** | $x(0) = 0$mM (for all metabolites)<br>$e(0) = 0$mM (for all enzymes) |

Supplementary Table S8: P-Aminostyrene model kinetic parameters

| Parameter | Symbol | Value | Units |
|---|---|---|---|
| Chorismate production rate | $V_{\text{chorismate}}$ | 1100. | 1/s |
| Deaminase production rate | $V_{\text{deaminase}}$ | 10. | 1/s |
| p-AF Loss | L | 1.4E-5 | 1/s |
| mRNA degradation rate | M | 3E-3 | 1/s |
| Protein degradation rate | P | 2E-4 | 1/s |
| Protein folding rate | F | 20 | 1/s |
| Dilution rate | $\delta$ | 5.79E-4 | 1/s |
| DNA duplication rate | $\beta$ | 5.78E-4 | 1/s |
| Avogadro's number | $\text{N}_{\text{A}}$ | 6.02214E23 | N/A |
| Cell volume | $\text{Vol}_{\text{cell}}$ | 2.5E-15 | L |
| Metabolite-induced toxicity | $t_a$ | 5E-4 | N/A |
| Protein-induced toxicity | $t_p$ | 50 | N/A |
| Enzyme-induced toxicity | $t_l$ | 50 | N/A |
| Toxicity constant | $k_i$ | 5E-5 | M |
| Pap operon mRNA length | $\text{L}_m$ | 3400 | nucleotides |
| Efflux pump mRNA length | $\text{L}_m$ | 2900 | nucleotides |
| LAAO mRNA length | $\text{L}_m$ | 1600 | nucleotides |
| Ribosome elongation rate | R | 20 | amino acids/s |
| Translation initiation rate | $\text{T}_{\text{init}}$ | 2E-1 | 1/s |
| Deaminase $k_{\text{cat}}$ | $k_{\text{cat}}$ | 5 | M/s |
| Deaminase $k_{\text{m}}$ | $k_{\text{m}}$ | 1E-6 | M |
| papA $k_{\text{cat}}$ | $k_{\text{cat}}$ | 0.2975 | M/s |
| papA $k_{\text{m}}$ | $k_{\text{m}}$ | 0.056 | M |
| papB $k_{\text{cat}}$ | $k_{\text{cat}}$ | 39 | M/s |
| papB $k_{\text{m}}$ | $k_{\text{m}}$ | 0.38 | M |
| papC $k_{\text{cat}}$ | $k_{\text{cat}}$ | 20.44 | M/s |
| papC $k_{\text{m}}$ | $k_{\text{m}}$ | 0.555 | M |
| LAAO $k_{\text{cat}}$ | $k_{\text{cat}}$ | 1.29 | M/s |
| LAAO $k_{\text{m}}$ | $k_{\text{m}}$ | 10.82 | M |
| Efflux pump rate | $k_{\text{m}}$ | 275 | M |

### S6 Hyperparameter tuning

The tree-structured Parzen estimator method (TPE) implemented in the Hyperopt package has a hyperparameter $\gamma$ which controls the balance of exploration and exploitation. TPE splits all samples into two distributions, a "good" and a "bad" distribution and selects the next point from the Expected Improvement computed over only the "good" samples. The parameter $\gamma$ is the fraction of points incorporated into the "good" distribution. The default value of $\gamma$ in the Hyperopt package is $\gamma = 15$, which corresponds to 15% of samples being partitioned into the "good" sample distribution and 85% in the "bad" distribution. Future samples are then drawn at the maximum of the "good" distribution. Higher values of $\gamma$ correspond to increased exploration of the loss landscape, whereas lower values of $\gamma$ mean increased exploitation towards local minima.

We found that regardless of the $\gamma$ value chosen, the distribution of architectures explored was similar (see Figure S1). To give a single scalar metric, we computed the fraction of samples in each quarter taken from the majority architecture, or the architecture most commonly sampled across the optimisation. This fraction was computed as

$$\frac{\text{majority architecture samples}}{\text{optimization iterations}}, \tag{S27}$$

for each quarter of the optimization trace. In the case of the glucaric acid model, the majority architecture is dual control. The majority architecture fraction rose with each quarter as the BayesOpt routine increasingly focused on low-loss dual control parameter value combinations; however, there was no statistical difference between the different gamma values chosen. These results show that the default $\gamma$ value is sufficient as TPE is relatively insensitive to hyperparameter tuning. This feature of TPE makes it quick to implement without significant application-specific tuning but may cause it to fail to adapt to certain specific cases. However, we did not observe TPE fail to converge in any of the systems under study.

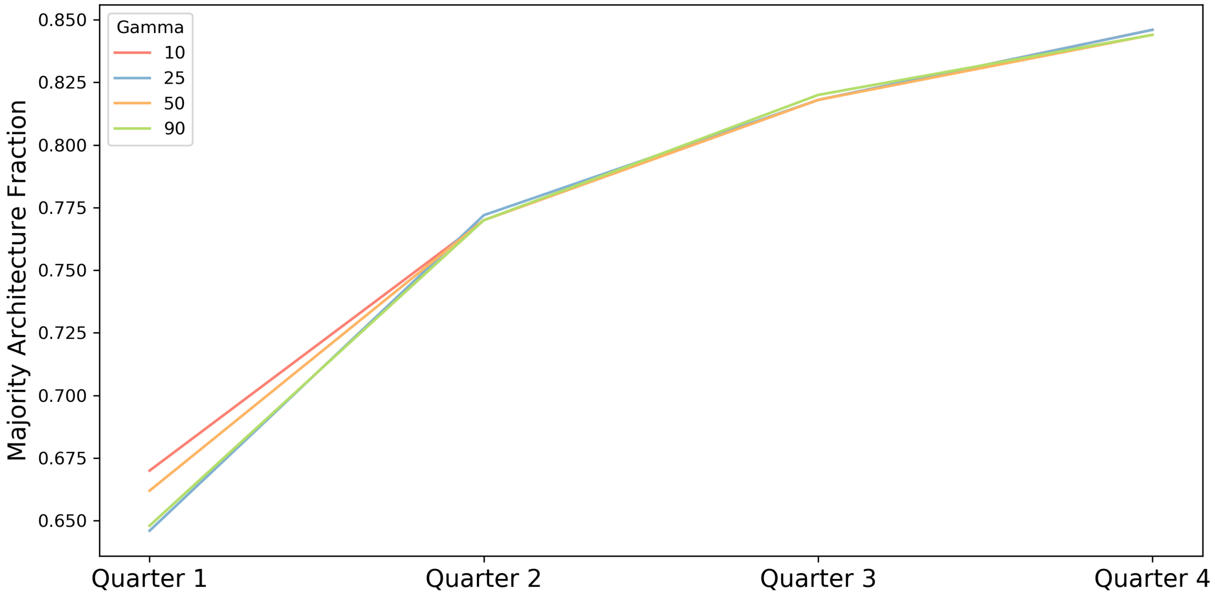


Supplementary Figure S1: **Hyperparameter Tuning** The TPE hyperparameter $\gamma$ was tuned by cloning the Hyperopt package repository and manually changing the value in the source code. Hyperparameter values of $\gamma = 10$, $\gamma = 25$, $\gamma = 50$, and $\gamma = 90$ were considered and a single optimisation of the glucaric acid pathway was run for each value. The total optimisations were split into four quarters by iteration and the percentage of sample drawn from the majority architecture (in this case, dual control).
